# Catalytic potential and disturbance rejection of glycolytic kinases in the human red blood cell

**DOI:** 10.1101/227058

**Authors:** James T. Yurkovich, Miguel A. Alcantar, Zachary B. Haiman, Bernhard O. Palsson

**Affiliations:** Department of Bioengineering, University of California, San Diego, La Jolla, CA; Bioinformatics and Systems Biology Program, University of California, San Diego, La Jolla, CA; Department of Pediatrics, University of California, San Diego, La Jolla, CA

## Abstract

The allosteric regulation of metabolic enzymes plays a key role in controlling the flux through metabolic pathways. The activity of such enzymes is traditionally described by allosteric rate laws in complex kinetic models of metabolic network function. As an alternative, we describe the fraction of the regulated enzyme that is in an active form by developing a detailed reaction network of all known ligand binding events to the enzyme. This fraction is the fundamental result of metabolic regulation as it represents the “tug of war” among the various regulators and substrates that determine the utilization of the enzyme. The active fraction corresponds to the utilization of the catalytic potential of the enzyme. Using well developed kinetic models of human red blood cell (RBC) glycolysis, we characterize the catalytic potential of its three key kinases: hexokinase (HEX), phosphofructokinase (PFK), and pyruvate kinase (PYK). We then compute their time-dependent interacting catalytic potentials. We show how detailed kinetic models of the management of the catalytic potential of the three kinases elucidates disturbance rejection capabilities of glycolysis. Further, we examine the sensitivity of the catalytic potential through an examination of existing personalized RBC models, providing a physiologically-meaningful sampling of the feasible parameter space. The graphical representation of the dynamic interactions of the individual kinase catalytic potential adjustment provides an easy way to understand how a robust homeostatic state is maintained through interacting allosteric regulatory mechanisms.

## Introduction

The human red blood cell (RBC) has historically been the target of complex kinetic model building of its metabolism due to its relative simplicity and the vast amounts of data and information available on its biochemistry and physiology. RBCs lack cellular compartments (e.g., nuclei, mitochondria) [1] and therefore certain cellular functions, such as transcriptional and translational regulation and the ability to use oxidative phosphorylation to produce energy [2]. As a result, glycolysis is the primary source of energy generation for the RBC, a pathway that undergoes allosteric regulation at major control points. Glycolytic ATP production is thus largely directly dependent upon the rate of ATP utilization.

Mathematical models have been used to study the dynamics of RBC metabolism since the 1970s [3]. Constraint-based modeling methods have been used to explore the mechanisms underlying cellular metabolism [4], [5], and specialized methods have been developed that allow for the study of system dynamics [6]–[8]. Kinetic models represent an approach that has the potential to truly capture the temporal dynamics at small time scales [9]. The first whole-cell kinetic model of RBC metabolism was published in the late 1980s [10]–[13], with other such models produced since then [14], [15]. More recently, enzyme modules have been introduced and used to explicitly model detailed binding events of ligands involved in allosteric regulation as an alternative to the traditional use of allosteric rate laws [16]. Such a fine-grained view of the activity and state of a regulated enzyme opens up many new possibilities in understanding the metabolic regulation that results from complex interactions of regulatory signals, as well as a way to explicitly represent biological data types.

Historically, the primary way to visualize the output from a kinetic model is to plot the time profiles of individual metabolite concentrations and enzymatic reaction rates. With the formulation of enzyme module models, we need to explore different ways of visualizing network dynamics because a different point of view brings about new insights, like Bode plots [17] or root loci [18] did in the early days of classical control theory. Enzyme modules allow for the explicit computation of the fraction of the regulatory enzyme in an active state that generates the reaction flux. Similar to how a controlled valve regulates water flow, the collective action of all the ligands binding to the enzyme—through the computation of the active enzyme fraction—fundamentally represent its regulation.

In this study, we use enzyme modules to model hexokinase (HEX), phosphofructokinase (PFK), and pyruvate kinase (PYK), the three major regulation points in RBC glycolytic energy generation. We compute the catalytic potential of these kinases as a measure of an enzyme’s capacity to influence the rest of the network, using the enzyme modules individually. We analyze the response of each enzyme module to perturbations in ATP utilization, simulating the impact of various physiological stresses on the RBC [19]–[21]. We then integrate all three enzyme modules into a single model of glycolysis and show that increasing the amount of regulation improves the disturbance rejection capabilities of the system to such perturbations. Finally, we elucidate how a graphical representation of the three kinase catalytic potentials leads to an insightful way to visualize the state of RBC glycolysis.

## Results

### Phosphofructokinase

PFK, often called the “pacemaker” of glycolysis [22], plays a major role in determining glycolytic flux. PFK converts fructose 6-phosphate (F6P) into fructose 1,6-bisphosphate (FDP). Here, we use a simple mechanism (Fig 1A): PFK must first bind to ATP, forming a complex that then binds F6P and converts it to FDP, producing ADP in the process. The four binding sites operate independently, i.e. they do not “cooperate.” The catalytic activity of PFK is controlled through allosteric regulation by AMP and ATP (Fig 1). AMP and ATP bind to an allosteric site, distal to the catalytic site, inducing a conformational change that modulates the activity of PFK.

**Fig. 1.**
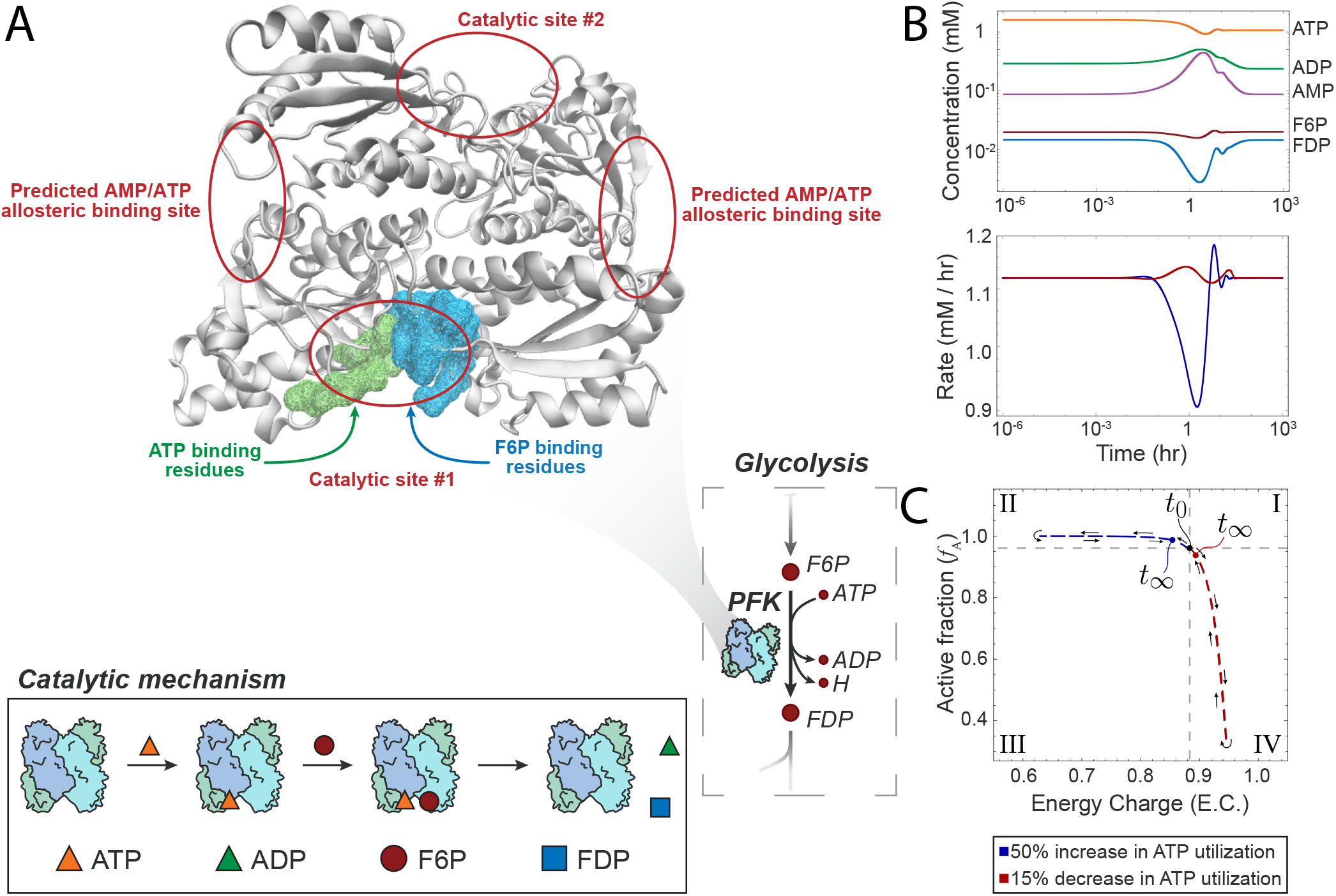
Overview of PFK mechanism and simulation results for PFK module in glycolysis only. (A) The structure of one of two PFK homomers along with the catalytic mechanism. The predicted allosteric binding sites for AMP/ATP are highlighted. (B) Concentration and reaction rate profiles for PFK regulatory module in glycolysis. The concentration profiles shown are for a 50 % increase in ATP utilization. (C) Phase portrait showing the catalytic potential of PFK. Two perturbations are shown: (1) a 50 % increase in ATP utilization, and (2) a 15 % decrease in ATP utilization. Roman numerals indicate comparisons with the steady-state: (I) more enzyme in active form and higher energy charge; (II) more enzyme in active form and lower energy; (III) more enzyme in inactive form and lower energy charge; and (IV) more enzyme in inactive form and higher energy charge.

For an enzyme allosterically regulated through effector molecules, we can define a quantity that relates the amount of enzyme in the active form to the total amount of enzyme. This catalytically active fraction (*f*_A_) is given by

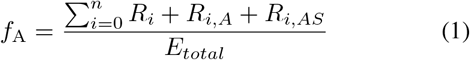

where *n* is the number of enzymatic binding sites, *R_i_* is the unbound enzyme in the active state (i.e., not bound to inhibitors), *R*_*i*,A_ is the enzyme bound to the cofactor, *R*_*i*,AS_ is the enzyme bound to the substrate and cofactor, and *E*_total_ is the total amount of enzyme. The subscript *i* represents the amount of activators bound to allosteric sites; for tetrameric structures like PFK and PYK, *i* ranges between 0 and 4 [23], [24].

In order to characterize the system response to perturbations, we modulated the ATP utilization by adjusting the rate of reaction for ATP. We modeled two perturbations that have been previously observed to fall within a physiologically feasible response: a 50% increase and a 15% decrease in ATP utilization [19]–[21]. We modeled these perturbations by modulating the rate of reaction for the hydrolysis of ATP (see Methods for full details). Increasing this rate decreases the amount of available ATP and ADP, resulting in a decrease in the rate of PFK (Fig 1B). Conversely, lowering this value increases the rate of the PFK reaction. For both perturbations, the final rate value eventually returns to the same as the unperturbed system. We were interested in characterizing the enzymatic response to these energetic perturbations. The energetic state of a cell can be measured using the energy charge [25], which relates the amount high energy bonds available in the adenosine phosphate pool. The energy charge is given by:

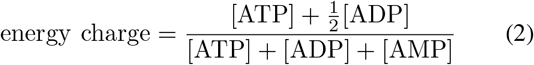

where [AMP], [ADP], and [ATP] represent the concentrations of those respective metabolites. To evaluate how the regulatory state of an enzyme is related to the energetic state of the system, we define the ratio of active to total enzyme as a function of the energy charge. We term this ratio the “catalytic potential” of an enzyme because it provides a representation of an enzyme’s affect on the rest of the system and its ability to maintain the homeostatic state.

We characterized our model of glycolysis with an enzyme module detailing the regulation of PFK, observing that PFK corrects for deviations in the energy charge by altering the amount of enzyme in the relaxed state (Fig 1C). We observed an inverse relationship between the energy charge and the catalytically active (relaxed) enzyme fraction which shows that an increase in ATP would inhibit PFK activity by shifting more enzyme into the catalytically inactive (tense) state.

### Hexokinase and pyruvate kinase

Having used an enzyme module to explicitly model the regulation of PFK, our goal was to expand our model to include the other glycolytic kinases (HEX and PYR). We constructed enzyme modules for both enzymes using mechanisms that allowed the substrate to bind cofactors in any order; hemoglobin was added to the model for HEX because it is necessary to model regulatory effects (see Methods for full details). We validated each module individually by performing the same ATP utilization perturbation (i.e., 50% increase and 15% decrease). The catalytic potential observed for the HEX module was in agreement with previously observed experimental evidence [26]. The PYK module exhibited a direct relationship between *f*_A_ and energy charge, conflicting with the inverse relationship previously observed *in vitro* [26]. There are several factors that could account for this discrepancy that focus on the scale and environmental factors of our model in comparison to the literature. The networks used in previous studies were on a much smaller scale than our network, negating the influence of other enzymes on PYK activity. Additionally, these assays did not contain FDP, which is a known activator of PYK. In our model, increasing the energy charge led to an initial increase in FDP concentration, which corresponded to an increase in the amount of PYK in the catalytically active form.

### Full kinase regulatory model (FKRM)

Once we had validated each kinase module individually, we built an expanded model of glycolysis that included all three enzyme modules and hemoglobin; we will refer to this model as the “full kinase regulatory model” (FKRM). We first calculated the concentration profiles for PFK (Fig 2A), observing higher FDP levels than with just the PFK module (Fig 1B). This effect is likely due to the inclusion of the PYK module, in which FDP is an allosteric activator. We also examined the rate profile of PFK (Fig 2B), which was inverted compared to that of the PFK model only (Fig 1B). This inversion arose from the addition of the HEX module, demonstrating the interplay among the various enzymes within a network.

**Fig. 2.**
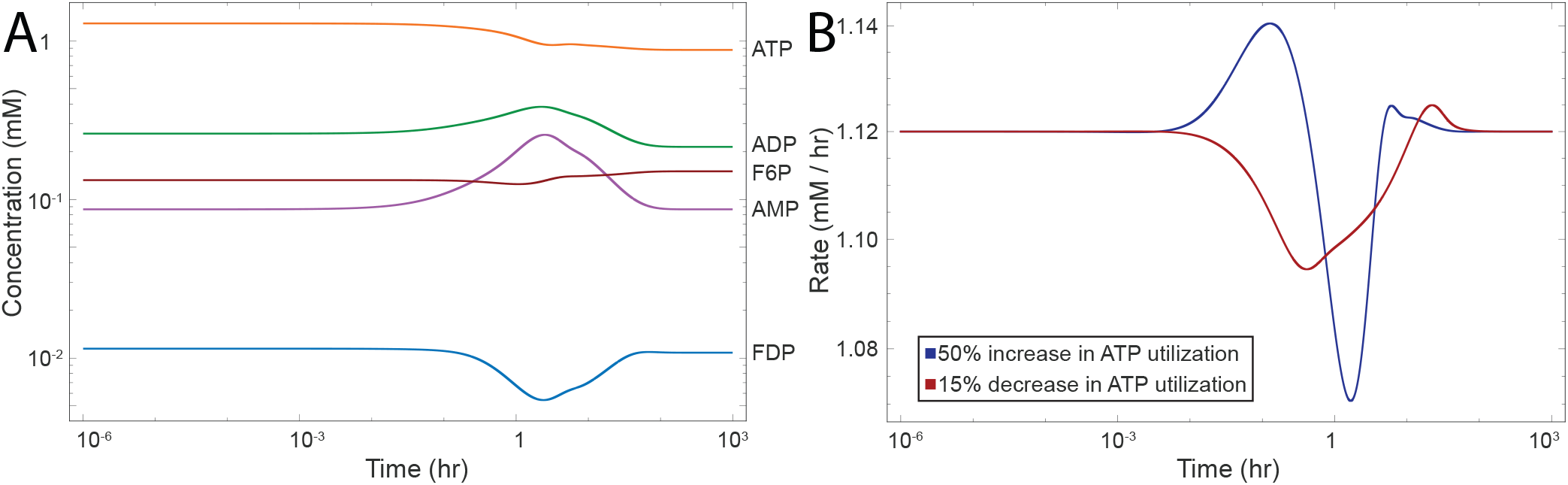
Classical representation of PFK simulation results for full kinase regulatory model. (A) Concentration time profiles shown for a 50% increase in ATP utilization. (B) Reaction rate time profiles for the reaction rate of PFK.

In order to better capture this interplay among enzymes, we constructed phase portraits for the fraction of catalytically active enzyme (*f*_A_) for each combination of enzyme modules in the FKRM (Fig 3A). These phase portraits show that as a greater fraction of PFK entered a more catalytically active state, a greater fraction of HEX become catalytically inactive; a similar behavior was observed for the PFK-PYK pair. We observed that HEX and PYK moved in tandem, with both enzymes moving into catalytically active or inactive states together. This behavior is likely due to the fact that these enzymes represent the boundaries of the system and therefore are linked in order to maintain system stability. Finally, we constructed catalytic potential plots for each enzyme module in the FKRM (Fig 3B). We observed that HEX and PYK exhibited primarily direct relationships between energy charge and the fraction of catalytically active enzyme, while we observed an inverse relationship between these two quantities for PFK. This inverse relationship observed for PFK recapitulated the previously reported relationship between catalytically active enzyme fraction and energy charge [27].

**Fig. 3.**
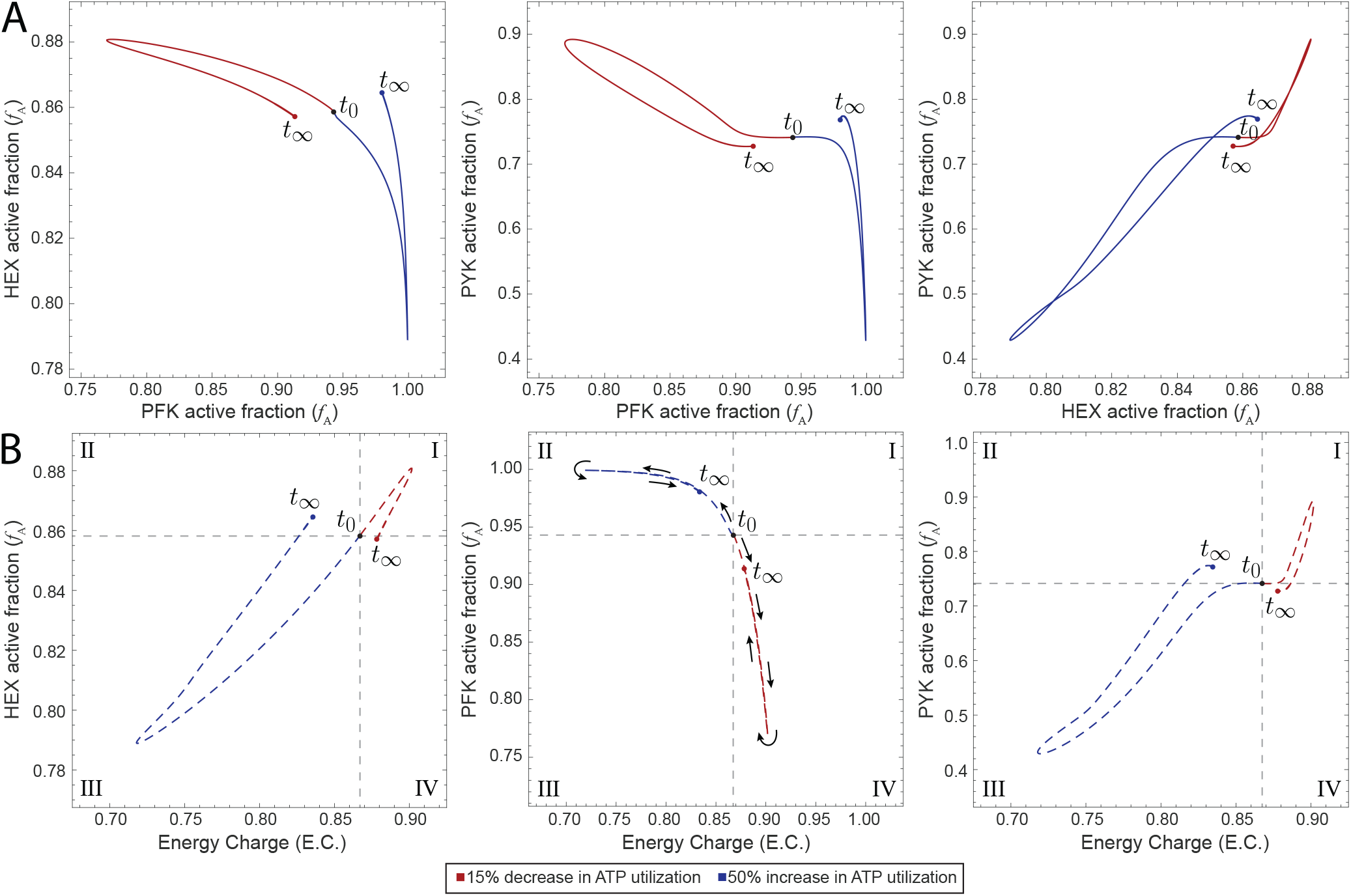
Characterization of full kinase regulatory model. (A) Phase portraits displaying pairwise relationships of the catalytic potentials of two kinases. (B) Catalytic potential plots for each of the enzyme modules in the full kinase regulatory model. Roman numerals indicate comparisons with the steady-state: (I) more enzyme in active form and higher energy charge; (II) more enzyme in active form and lower energy; (III) more enzyme in inactive form and lower energy charge; and (IV) more enzyme in inactive form and higher energy charge.

### Disturbance rejection capabilities

The inclusion of feedback and regulation mechanisms improves the disturbance rejection capabilities of a system [28]. For biological systems, these regulatory effects help organisms maintain the homeostatic state. Having characterized our individual enzyme modules and the FKRM, our next goal was to investigate (1) the capacity for each of these models to maintain the homeostatic state and (2) how examining the catalytic potential helped elucidate these behaviors. We modeled a 50% increase in ATP utilization and calculated the total ATP flux in the network (i.e., total flux through ATP-producing reactions minus total flux through ATP-consuming reactions) for each of the models we constructed (Fig 4). All systems were able to retain to a stable homeostatic state following the perturbation (Fig 4A). We calculated the sum of squared error (SSE) for each model in order to quantify the disturbance rejection capabilities of each model (Fig 4A). As expected, the models with little or no regulation performed the worst, while increased regulation generally lowered the SSE. The base glycolytic model with the PYK module performed the worst, while the model containing the PFK and HEX modules with hemoglobin performed the best. The final steady-state values for the energy charge differed with the inclusion of hemoglobin in the model, although the magnitude of these differences was small (Fig 4B).

**Fig. 4.**
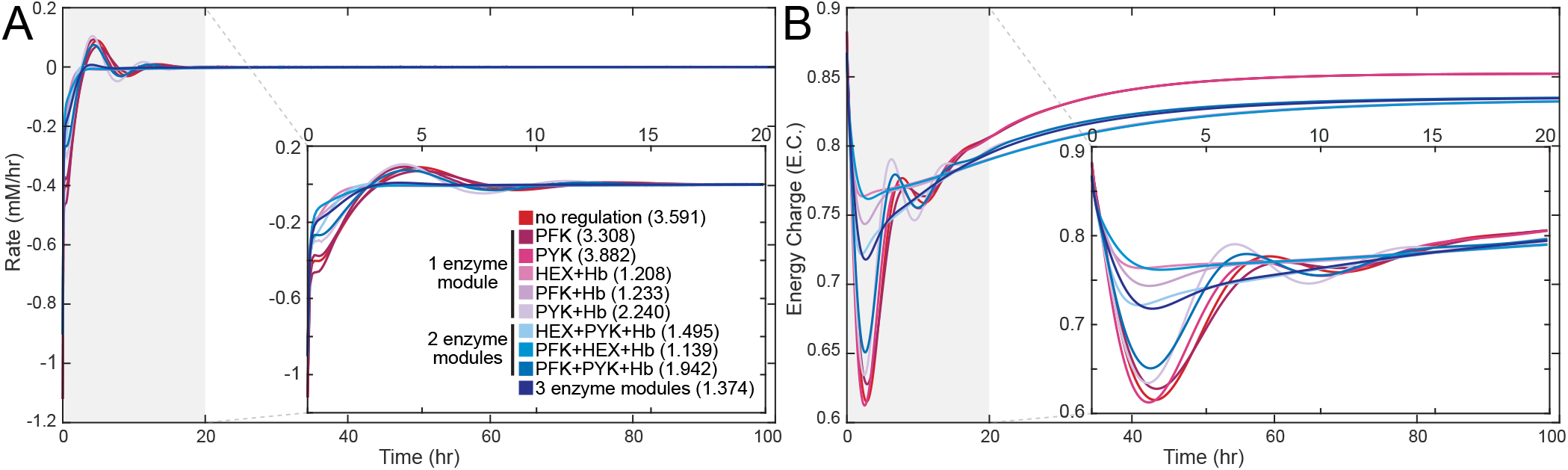
Disturbance rejection capabilities of glycolytic models with varying amounts of regulation. (A) The net rate of ATP usage (i.e., total flux through ATP-producing reactions minus total flux through ATP-consuming reactions) is shown as a function of time; the inset zooms in on the 0 to 20 hour time range. The number in parentheses represents the SSE for each model, quantifying the total deviation of the output from the setpoint. (B) The energy charge is shown as a function of time; the inset zooms in on the 0 to 20 hour time range.

Rather than perform a sensitivity analysis by randomly perturbing parameters in the kinetic equations, we instead chose to examine the disturbance rejection capabilities of personalized models constructed from measurements obtained from multiple individuals; these models represent a more realistic sampling of physiologically-feasible model parameters. We constructed an individual model using glycolytic metabolite concentrations and equilibrium constants for nine individuals from Bordbar et al. [15]. We performed our sensitivity analysis using the model that includes a PFK module and hemoglobin due to simplicity for numerical simulation. The general qualitative trend for the catalytic potential plot was similar to the one using literature values (Fig 1C and Fig 3B), but initial *f*_A_ values were significantly lower in the personalized models (Fig 5A,B). In particular, the amount of active PFK for each individual reached a saturation point that was higher than the initial steady-state value in order to compensate for the increase in ATP utilization before returning to a final steady-state value. While we observe that there is little difference among the rate profiles (Fig 5C), we observe much greater differences in the catalytic potential plots (Fig 5A,C) and energy charge profiles (Fig 5D). Notably, the model for Individual #1 exhibited a much different response than the other eight personalized models (Fig 5A,B,D). Upon further examination, we determined that this difference stemmed from the fact that the rate constants for the binding of ATP and F6P to PFK were outliers with over 99% confidence according to the Dixon’s Q test (see Methods for full details); these were the only rate constants that were deemed to be outliers out of all enzymatic reactions.

**Fig. 5.**
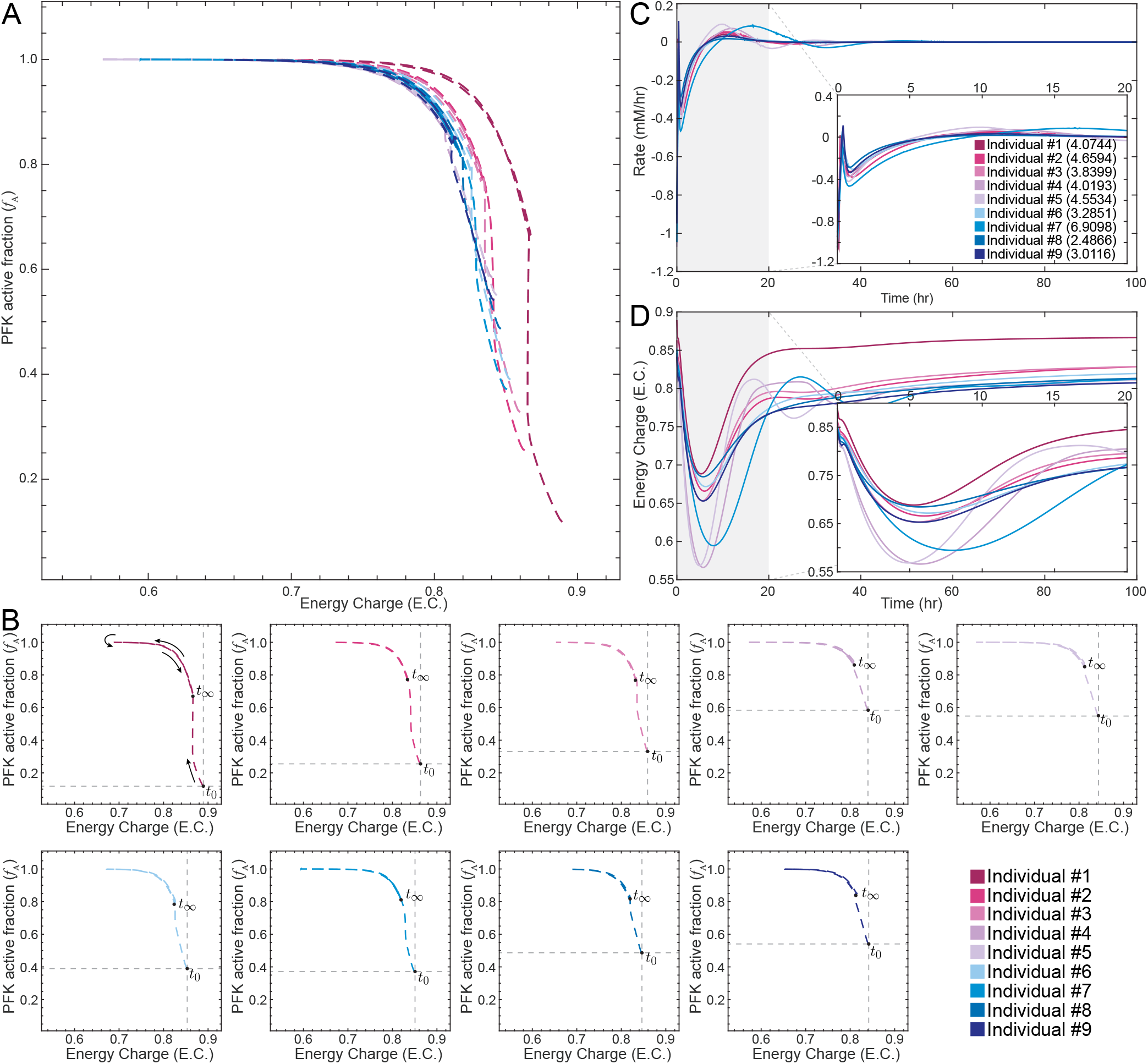
Disturbance rejection capabilities of personalized glycolytic models with a PFK module and hemoglobin. (A) Superimposed catalytic potential plots for all personalized models. (B) Catalytic potential plots for each individual;the intersection of the gray lines denotes the initial steady-state value at time zero and helps show the differences among the population. (C) The net rate of ATP usage (i.e., total flux through ATP-producing reactions minus total flux through ATP-consuming reactions) is shown as a function of time; the inset zooms in on the 0 to 20 hour time range. The number in parentheses represents the SSE for each model, quantifying the total deviation of the output from the setpoint. (D) The energy charge is shown as a function of time; the inset zooms in on the 0 to 20 hour time range.

## Discussion

The ability to mechanistically model cellular metabolism allows for the construction of predictive physiological models. However, the mechanistic results obtained from time-course plots can complicate the interpretation and analysis of systems-wide responses to relevant perturbations. To help provide a better method of elucidating this behavior, we built modularized glycolytic models with enzymes serving as regulators. These models were then validated against existing empirical data to understand the relationship between the catalytically active enzyme fraction and energy charge—the catalytic potential of an enzyme. Visualizing the catalytic potential allowed for the analysis of important systems behaviors. The results presented here have several primary implications.

First, we have studied glycolysis from a perspective in which enzymes are regulators. Individual kinases serve as tuning dials for the system. Adjusting these dials changes the response of the system, as demonstrated by examining individual parameterization of personalized models (Fig 5). Through an examination of the catalytic potential of PFK, we were able to gain insight into how the regulator within a model is tuned in different individuals in order to maintain homeostasis (Fig 5A,B,D), a behavior that was not discernible through more typical metrics like rates of reaction (Fig 5C). While the present results are limited by the scope of the model (i.e., only glycolysis), expanding this framework to larger-scale models of RBC metabolism could provide similarly interesting results.

Second, the disturbance rejection capabilities of the models improved with the incorporation of additional regulatory mechanisms (Fig 4A). We simulated physiologically-relevant perturbations, observing that systems with regulation are improved over those with less regulation (i.e., fewer modules) as shown by quantifying the total deviation of the model output from the setpoint (i.e., the SSE). It is notable that models with hemoglobin and either HEX or PFK performed well despite not accounting for all regulatory mechanisms, indicating that kinetic models that do not account for the regulation in these important steps in glycolysis fail to capture important behaviors that affect the rest of the network. It is likely that the vast improvement of those two models over the model with PYK and hemoglobin is due to the fact that PYK is one of the last steps in glycolysis and therefore has a smaller impact on the rest of the system. We further investigated the disturbance rejection capabilities of the PFK and hemoglobin model through a sensitivity analysis that used physiologically-relevant parameterizations instead of randomly-distributed parameter sets. This analysis helped elucidate subtle differences among individuals that were accessible only by studying the systems-level effects of regulation.

Finally, we have shown that the catalytic potential is a metric that can provide additional insight into how metabolic networks maintain a homeostatic state following physiologically-relevant perturbations. Using a small-scale model that explicitly accounted for the regulatory mechanisms of the three glycolytic kinases, we investigated the interplay between these three enzymes. When we applied this metric to examine the response of personalized models to ATP utilization perturbations, we observed differences that were not apparent simply from the rate profile. Upon further investigation, we were able to hypothesize that the catalytic potential for that individual was different than the others due to differences in the binding of ATP and F6P to PFK. Thus, the catalytic potential helped provide insight into how subtle differences among individuals can lead to differing systemic responses to perturbations that push the system away from the homeostatic state.

Red blood cells are networks consisting of well-studied metabolic pathways and their associated metabolites. However, it is often difficult to examine individual enzymes *in vivo* without using small scale assays [26], [27], [29], [30]. These assays are not comprehensive and, as a result, may not provide an accurate depiction of the interplay between multiple regulatory enzymes in a network like glycolysis. New methods of visualizing this behavior—such as the catalytic potential plot introduced here—can lead to new insights and discoveries. Viewing enzymes as regulators through which we can tune the system response opens the door for us to investigate what the optimal state might be and how that state helps maintain homeostasis.

## Methods

All calculations were performed in Mathematica 11.1 [31]. Simulations were conducted using the Mass Action Stoichiometric Simulation (MASS) Toolbox kinetic modeling package (https://github.com/opencobra/MASS-Toolbox). Details for formulating a MASS model are found in Jamshidi et. al. [32]. All models used are available upon request.

### Glycolysis and the Rapoport-Luebering shunt

The base glycolysis network included all 10 glycolytic enzymes and lactate dehydrogenase. Reaction rates were defined using mass action kinetics, representing enzyme catalysis as a single step. These simplified reactions were systematically replaced with enzyme modules following the procedure outlined by Du et al. [16]. Additionally, a phosphate exchange reaction was incorporated into the glycolytic network utilizing parameters obtained from Prankerd et al. [33]. Similarly, the Rapoport-Luebering shunt was included in some models to account for the presence of hemoglobin, whose binding to oxygen is regulated by 2,3-diphosphoglycerate (2,3-DPG). Incorporation of this shunt was accompanied by parameter changes as previously described [34].

### Enzyme module construction

Regulation was manually incorporated into the enzyme reactions. Initial conditions from the glycolysis and hemoglobin MASS toolbox example data were used in conjunction with equilibrium constants which were obtained from from [35], [36]. These values were subsequently utilized to solve for new kinetic parameters. This procedure (outlined in [34]) adheres to the formula:

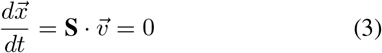

where 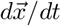 is the concentration rate of change with respect to time for metabolites, **S** is the stoichiometric matrix, and 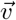 is a vector containing reaction fluxes.

We constructed a total of ten different models with varying amounts of regulation, spanning from the base glycolytic model with no enzyme modules (and therefore no regulation) to the FKRM with three enzyme modules and the Rapaport-Luebering shunt. The remaining models represented each combination of the three kinase modules. Enzyme module incorporation was accompanied by the deletion of the original single-step reaction in order to avoid redundant reactions. Stability for all systems was verified by simulating the network and ensuring that a steady-state point was found for all metabolites.

#### Hexokinase (HEX)

HEX (EC 2.7.1.1) was modeled as a monomer to account for the fact that it contains only one active catalytic site. The previously specified mechanism was chosen to match that used by [16] because all kinetic parameters were obtained from this source. A hemoglobin module is necessary to include when the HEX module is included because it affects the level of 2,3 DPG, which serves as a regulatory molecule for HEX. The HEX module consisted of the following chemical reactions:

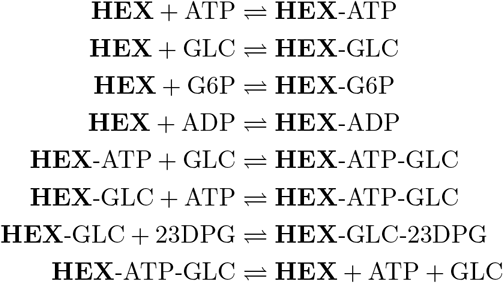

where the bold text represents the enzyme and dashes show bound species.

#### Phosphofructokinase (PFK)

PFK (EC 2.7.1.11) was modeled as a homotetramer to account for its four catalytic and allosteric binding sites [37]. The previously specified mechanism was chosen to match that used by [16] because all kinetic parameters were obtained from this source. The PFK module consisted of the following chemical reactions:

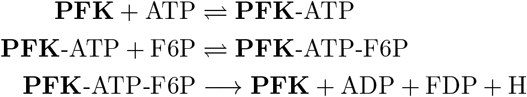

where the bold text represents the enzyme and dashes show bound species. Additional reactions were included to account for the conversion between the tight and relaxed state, as well as the effector molecule binding.

#### Pyruvate kinase (PYK)

PYK (EC 2.7.1.40) was modeled to include allosteric regulation. Additional reactions were also included to account for the equilibration of both enzymes between the relaxed (R) and tense (T) state [24]. Additionally, PYK was modeled as a tetramer to account for the four catalytic and allosteric sites on each enzyme. Dissociation constants were obtained from [12] and rate constants were solved using equation 3. The PYK module consisted of the following chemical reactions:

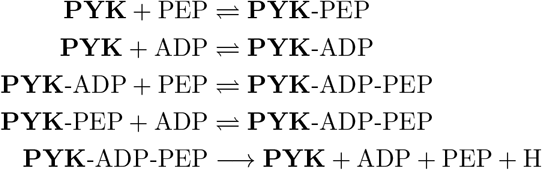

where the bold text represents the enzyme and dashes show bound species.

### Personalized models

Personalized models were constructed by replacing all intracellular glycolytic metabolite concentrations and equilibrium constants with values reported by Bordbar et al. [15]. New pseudo-elementary rate constant (PERC) values were calculated using the personalized concentration data. The Rapoport-Luebering shunt was added to the RBC network and PFK enzyme modules were created for all individuals using the resulting concentration values after the addition of the Rapoport-Luebering pathway. Due to numerical issues when attempting to simulate, we only used 9/24 of the models available in [15]. Individuals #1-9 in our study correspond to individuals 2, 4, 5, 6, 7, 8, 10, 16, and 18, respectively, from [15].

To identify outliers within the reaction PERCs compared with the other personalized models, we performed a Dixon’s *Q* test [38]:

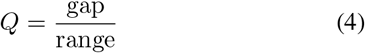

where the gap is the absolute difference between the point in question and the nearest value, and the range is the range of all values. For a set with nine samples, we can be 99% confident that a point is an outlier if the *Q* value is greater than 0.598; the *Q* values for the ATP and F6P binding steps had *Q* values of 0.84257 and 0.73164, respectively.

### System analysis

Rate pools for enzymes were defined as the rate of at which enzyme produced product. This was accomplished by defining a pool from the product’s ODE consisting solely of the terms contributing to product formation. In other words:

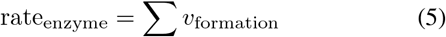

where *v*_formation_ represents the forward rate of the enzyme reaction and possesses units of mmol/L · s. Defining the rate pools in this manner neglected effects of reversible reactions contributing to the formation of product. Thus, this pool quantified the actual catalytic activity of the enzyme of interest.

### Simulating ATP utilization perturbations

In order to mimic a physiologically-relevant perturbation away from the homeostatic state, we simulated a 50% increase in ATP utilization and a 15% decrease in ATP utilization [19]-[21]. Changes in ATP utilization were applied by changing the rate (*k*_ATP_) associated with ATP hydrolysis:

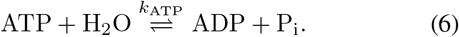

We calculated the sum of squared error (SSE) for each model in order to quantify the total deviation of the output from its setpoint, which is zero. The resulting quantity (i.e., the SSE) is compared between models, with a smaller value indicating better disturbance rejection capabilities.

## Acknowledgments

The authors gratefully acknowledge Nathan Mih, Laurence Yang, and Bin Du for valuable discussions. The authors would also like to thank the UC San Diego Academic Enrichment Programs Office and the Genentech Foundation Scholars Program.

## Author Contributions

JTY and BOP conceived the study. JTY, MAA, and ZBH performed analysis. JTY, MAA, and BOP wrote the manuscript. All authors read and approved the final manuscript.

## References

1 Zhang ZW, Cheng J, Xu F, Chen YE, Du JB, Yuan M, et al. Red blood cell extrudes nucleus and mitochondria against oxidative stress. IUBMB Life. 2011;63(7):560–565. doi:10.1002/iub.490.

2 Yoshida T, Shevkoplyas SS. Anaerobic storage of red blood cells. Blood Transfusion. 2010;8(4):220.

3 Pujo-Menjouet L. Blood Cell Dynamics: Half of a Century of Modelling. Mathematical Modelling of Natural Phenomena. 2016;11(1):92–115. doi:10.1051/mmnp/201611106.

4 Lewis NE, Nagarajan H, Palsson BO. Constraining the metabolic genotype–phenotype relationship using a phylogeny of in silico methods. Nature Reviews Microbiology. 2012;doi:10.1038/nrmicro2737.

5 Yurkovich JT, Palsson BO. Solving Puzzles With Missing Pieces: The Power of Systems Biology. Proceedings of the IEEE. 2016;104(1):2–7. doi:10.1109/jproc.2015.2505338.

6 Mahadevan R, Edwards JS, Doyle FJ. Dynamic flux balance analysis of diauxic growth in Escherichia coli. Biophysical journal. 2002;83(3):1331–1340.

7 Waldherr S, Oyarzún DA, Bockmayr A. Dynamic optimization of metabolic networks coupled with gene expression. Journal of theoretical biology. 2015;365:469–485.

8 Bordbar A, Yurkovich JT, Paglia G, Rolfsson O, Sigurjónsson Ólafur E, Palsson BO. Elucidating dynamic metabolic physiology through network integration of quantitative time-course metabolomics. Scientific Reports. 2017;7:46249.

9 Jamshidi N, Palsson B. Formulating genome-scale kinetic models in the post-genome era. Mol Syst Biol. 2008;4:171.

10 Joshi A, Palsson BO. Metabolic dynamics in the human red cell. Part I–A comprehensive kinetic model. J Theor Biol. 1989;141(4):515–528.

11 Joshi A, Palsson BO. Metabolic dynamics in the human red cell. Part II–Interactions with the environment. J Theor Biol. 1989;141(4):529–545.

12 Joshi A, Palsson BO. Metabolic dynamics in the human red cell. Part III–Metabolic reaction rates. J Theor Biol. 1990;142(1):41–68.

13 Joshi A, Palsson BO. Metabolic dynamics in the human red cell. Part IV–Data prediction and some model computations. J Theor Biol. 1990;142(1):69–85.

14 Nakayama Y, Kinoshita A, Tomita M. Dynamic simulation of red blood cell metabolism and its application to the analysis of a pathological condition. Theor Biol Med Model. 2005;2:18.

15 Bordbar A, McCloskey D, Zielinski DC, Sonnenschein N, Jamshidi N, Palsson BO. Personalized Whole-Cell Kinetic Models of Metabolism for Discovery in Genomics and Pharmacodynamics. Cell Syst. 2015;1(4):283–292.

16 Du B, Zielinski DC, Kavvas ES, Dräger A, Tan J, Zhang Z, et al. Evaluation of rate law approximations in bottom-up kinetic models of metabolism. BMC Systems Biology. 2016;10(1). doi:10.1186/s12918-016-0283-2.

17 Bode HW. Variable equalizers. The Bell System Technical Journal. 1938;17(2):229–244.

18 Cavicehi TJ. Phase-root locus and relative stability. IEEE Control Systems Magazine. 1996;16(4):69–77.

19 Wan J, Ristenpart WD, Stone HA. Dynamics of shear-induced ATP release from red blood cells. Proceedings of the National Academy of Sciences. 2008;105(43):16432–16437.

20 Arciero JC, Carlson BE, Secomb TW. Theoretical model of metabolic blood flow regulation: roles of ATP release by red blood cells and conducted responses. AJP: Heart and Circulatory Physiology. 2008;295(4):H1562–H1571. doi:10.1152/ajpheart.00261.2008.

21 Abraham EH, Salikhova AY, Hug EB. Critical ATP parameters associated with blood and mammalian cells: Relevant measurement techniques. Drug Development Research. 2003;59(1):152–160. doi:10.1002/ddr.10194.

22 Berg JM, Tymoczko JL, Stryer L. Biochemistry (Chapters 1-34). W. H. Freeman; 2002. Available from: https://www.amazon.com/Biochemistry-Chapters-1-34-Jeremy-Berg/dp/0716730510?SubscriptionId=0JYN1NVW651KCA56C102&tag=techkie-20&linkCode=xm2&camp=2025&creative=165953&creativeASIN=0716730510.

23 Schöneberg T, Kloos M, Brüser A, Kirchberger J, Sträter N. Structure and allosteric regulation of eukaryotic 6-phosphofructokinases. Biological Chemistry. 2013;394(8). doi:10.1515/hsz-2013-0130.

24 Zanella A, Fermo E, Bianchi P, Valentini G. Red cell pyruvate kinase deficiency: molecular and clinical aspects. British Journal of Haematology. 2005;130(1):11–25. doi:10.1111/j.1365-2141.2005.05527.x.

25 Atkinson DE, Walton GM. Adenosine triphosphate conservation in metabolic regulation rat liver citrate cleavage enzyme. Journal of Biological Chemistry. 1967;242(13):3239–3241.

26 Purich DL, Fromm HJ. Studies on factors influencing enzyme responses to adenylate energy charge. Journal of Biological Chemistry. 1972;247(1):249–255.

27 Shen L, Fall L, Walton GM, Atkinson DE. Interaction between energy charge and metabolite modulation in the regulation of enzymes of amphibolic sequences. Phosphofructokinase and pyruvate dehydrogenase. Biochemistry. 1968;7(11):4041–4045.

28 Zames G. Feedback and optimal sensitivity: Model reference transformations, multiplicative seminorms, and approximate inverses. IEEE Transactions on Automatic Control. 1981;26(2):301–320. doi:10.1109/tac.1981.1102603.

29 Liao CL, Atkinson DE. Regulation at the phosphoenolpyruvate branchpoint in Azotobacter vinelandii: pyruvate kinase. Journal of bacteriology. 1971;106(1):37–44.

30 Ling KH, Byrne WL, Lardy H. [38] Phosphohexokinase: Fructose-6-phosphate+ ATP→ Fructose-1, 6-diphosphate+ ADP. Methods in enzymology. 1955;1:306–310.

31 Wolfram Research Inc. Mathematica 11.1; 2017. Available from: http://www.wolfram.com.

32 Jamshidi N, Palsson BØ. Mass Action Stoichiometric Simulation Models: Incorporating Kinetics and Regulation into Stoichiometric Models. Biophysical Journal. 2010;98(2):175–185. doi:10.1016/j.bpj.2009.09.064.

33 Prankerd TAJ, Altman KI. A Study of the Metabolism of Phosphorus in Mammalian Red Cells. Biochemical Journal. 1954;58(4):622–633.

34 Palsson BO. Systems Biology: Simulation of Dynamic Network States. New York: Cambridge University Press; 2011.

35 Ponce J, Roth S, Harkness DR. Kinetic studies on the inhibition of glycolytic kinases of human erythrocytes by 2, 3-diphosphoglyceric acid. Biochimica et Biophysica Acta (BBA)-Enzymology. 1971;250(1):63–74.

36 Gerber G, Preissler H, Heinrich R, Rapoport SM. Hexokinase of Human Erythrocytes. Purification, Kinetic Model and Its Application to the Conditions in the Cell. European Journal of Biochemistry. 1974;45(1):39–52. doi:10.1111/j.1432-1033.1974.tb03527.x.

37 Hoggett JG, Kellett GL. Kinetics of the cooperative binding of glucose to dimeric yeast hexokinase PI. Biochemical journal. 1995;305(2):405–410.

38 Dean RB, Dixon WJ. Simplified Statistics for Small Numbers of Observations. Analytical Chemistry. 1951;23(4):636–638. doi:10.1021/ac60052a025.

